# Chemically-defined induction of a primitive endoderm and epiblast-like niche supports post-implantation progression from blastoids

**DOI:** 10.1101/510396

**Authors:** Erik J. Vrij, Yvonne S. Scholte op Reimer, Javier Frias Aldeguer, Isabel Misteli Guerreiro, Jop Kind, Bon-Kyoung Koo, Clemens A. van Blitterswijk, Nicolas C. Rivron

**Author notes:** Correspondence should be addressed to: Nicolas Rivron -, Erik Vrij.

## Abstract

The early mammalian conceptus (blastocyst) contains two supporting extraembryonic tissues - the trophectoderm and the primitive endoderm (PrE) - that encase and guide the epiblast (Epi) to eventually form the all body. Modifications of the conceptus exposed key genes regulating these tissues co-development. However, the combinations of signalling pathways underlying the interplay of PrE and Epi remains elusive. Stem cell-based models including embryoid bodies and blastoids can be generated in large numbers and subjected to high-content screens. Here, we use combinatorial screens of proteins, GPCR ligands and small molecules to rapidly (72 hours) and efficiently (80%) guide embryoid bodies to form a three-dimensional PrE-/Epiblast-like niche in chemically-defined conditions (gel-free, serum-free). This bipotent niche spontaneously progresses, without growth factors, to form a pro-amniotic cavity surrounded by a polarized Epi covered with parietal and visceral endoderm-like cells. In blastoids, these molecules enhance the ratio and number of Gata6+/Nanog+ cells and promote the survival, expansion and morphogenesis of a post-implantation-like Epi *in vitro*. Altogether, modelling early development in chemically-defined conditions delineates the pathways sufficient to form a functional PrE/Epiblast niche that fuels post-implantation development.

## Introduction

The early mammalian conceptus consists of three lineages: the pluripotent Epiblast (Epi) that forms the embryo proper, and the two extraembryonic lineages — trophoblast and primitive endoderm (PrE) — that contribute to the placenta and yolk sac, respectively^1,2^. In mice, the bifurcation between PrE and Epi cells is established in a seemingly sequential and synergistic manner between E3.25 and E4.5^3–6 7^, and marked by the timed expression of transcription factors Gata6, PDGFRα, Gata4, Sox17, and Sox7^2,8,9^. Experiments using *in vivo* and *in vitro* models suggested that this process is initiated by lineage priming^10^ exploiting polycombs^11^, chromatin modifiers^12^ and small-RNAs^13^ activities, along with the progression of gene regulatory networks^2^ and intercellular signalling circuitries (e.g. FGF/Mapk/Erk^5,10,14–21^ and Lif/Stat^4,22^, Bmp4/Smad4^23,24^ and Wnt/β-Catenin^25,26^ pathways). In the mouse blastocyst this initial cell fate choice is reinforced by a PrE/Epi cross-talk^8,27–31^, which progressively locks cell fates, promote their physical segregation, and the epithelization and lining of the PrE along the blastocoel cavity^32–34,35^ while adjusting the mutually allocated cell numbers^20,36–38^.

The resolution of a niche (E4.5) comprising two outer tissues (the polar trophoblast and the PrE) encasing the inner Epi acts as a checkpoint for developmental progression, coincides with the blastocyst implantation into the uterus, and leads to both a switch in Epi identity (pluripotent-formative-primed states)^39–42^ and the morphogenesis of a rosette^43,44^. The formation of a basement membrane, mainly by the trophoblast and PrE cells, along with differential molecular signalling (e.g. Nodal^45^ and Lif^46^) and mechanical properties^47^ are likely to support the sustainability and polarization of the Epi with centrally oriented apical sides. This polarization coincides with a switch of the core transcriptional network and epigenetic state of the Epi^39–42^ marked by a loss of *Nanog* and a gain of *Otx2*^*48,49*^, and with the induction, expansion and fusion of lumens in part^45^ *via* beta-integrin signalling^50^ and exocytosis of negatively charged sialomucins including Podocalyxin (Podxl) that repulse the apical cell membranes^43,46,51,52^. These processes resolve the primed Epi state and the pro-amniotic cavity, which altogether set the stage for post-implantation development.

The formation of PrE and derivatives has been studied in embryoid bodies (EBs) - aggregates of embryonic stem cells (ESCs) that reflect aspects of early embryonic development^53,54^. However, EBs are usually cultured in conditions that do not favour a precise control over cell number and confinement, nor the chemical nature of the medium. As a result, the PrE does not form efficiently and the culture parameters (e.g. the use of serum-containing medium) occlude underlying mechanisms. In contrast, the use of microsystems^55^ and chemically-defined medium^55,56^ opens possibilities to increase the control, throughput and screening capacities, thus better understand cell fate.

Here we run combinatorial screens of proteins, GPCR ligands and small molecules in a microwell array platform and in chemically-defined conditions. This pipeline establishes a cell culture model that rapidly and efficiently co-form PrE- and Epi-like cells amenable to in-depth investigation. Applying this novel model to test the functional co-development of embryonic and extraembryonic tissues^57^, we investigate how tissues synergies support the potential for viability, expandability and morphogenesis of the post-implantation epiblast.

## Results

### Naïve pluripotency enhances the ESCs potential for PrE differentiation

We used a high-content screening platform of non-adherent hydrogel microwells in 96-well-plates^58^ to reproducibly aggregate small, defined numbers of ESCs into EBs (Figure 1A). The number of ESCs seeded into microwells followed a Poisson distribution across the 430 microwells (7–12 cells per microwell), which aggregated within 24 hours (Figure 1B, **Figure S1)**. We quantified PrE differentiation *via* in situ imaging of a fluorescent reporter under the promoter for Pdgfrα (ESCs ^Pdgfrα-h2b-gfp/+^, **Figure S2**)^7,59^. EBs survived in serum-free B27N2 medium and leukemia inhibitory factor (Lif) but did not proliferate and formed only a few PrE cells (Yield of Pdgfrα+ EBs: 1%, Figure 1C, **Figure S3**). In contrast, the addition of serum induced proliferation and the appearance of Pdgfrα+ EBs (44%, Figure 1C, *p* < 0.001, ANOVA with Tukey’s multiple comparison test). Consistent with a previous report^18^, we observed that an initial 2D expansion in chemically-defined B27N2/2i/Lif medium^60^ enhanced the permittivity for PrE formation, as compared to an initial expansion in serum-containing medium (Figure 1C). Thus *naïve* ESCs have an enlarged ability to respond to PrE-inductive signals. We concluded that similar to the blastocyst cells^7,59^, PrE formation in EBs requires an initial permissive state, along with undefined signals regulating proliferation and differentiation.

**Figure 1:**
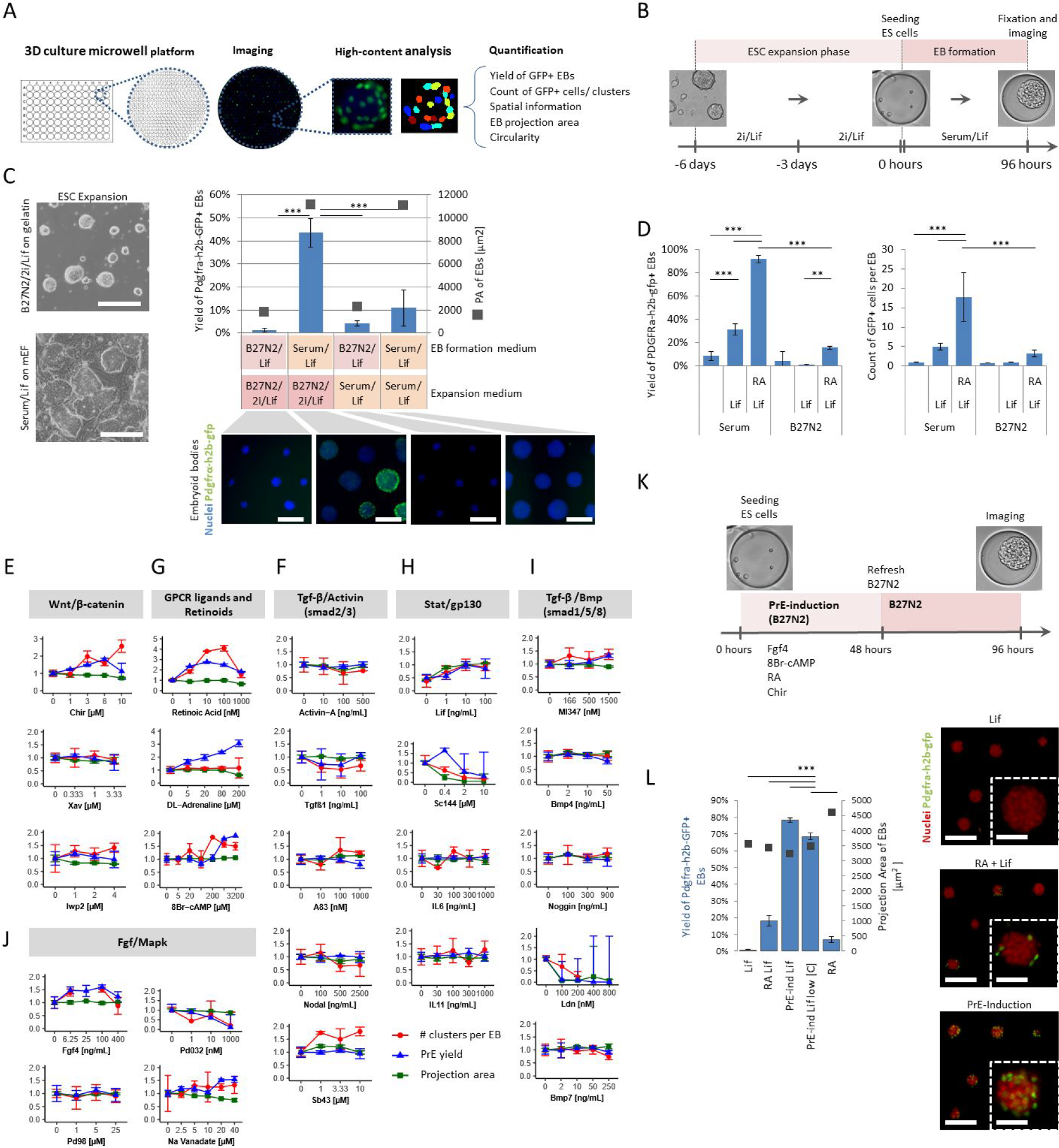
The initial naive state of ESCs and specific signalling pathways induce the efficient and rapid formation of PrE-/Epi-like niche *in vitro*. (A) High-content screening (HCS) methodology of 96-well plates imprinted with agarose microwell arrays (430 microwells per well) in which EBs are formed, cultured and imaged (each microwell capturing a single EB). (B) Schematic of experimental set-up including ESC expansion and EB-based primitive endoderm (PrE) differentiation. (C) (left) Bright-field images of ESCs expanded in B27N2/2i/Lif or serum/Lif on mouse embryonic fibroblasts (mEF). (right) Quantified yield of PrE-differentiation (Pdgfrα+, left axis) and size of EBs (projection area, right axis) derived from ESCs expanded in naïve (B27N2/2i/Lif) versus serum/Lif conditions. (bottom) Fluorescence images show the nuclei (blue) and Pdgfrα-h2b-gfp+ cells (green) within EBs formed in either B27N2/Lif from ESCs expanded in B27N2/2i/Lif (left), or serum/Lif from ESCs expanded in B27N2/2i/Lif (middle left), or B27N2/2i/Lif from ESCs expanded in serum/Lif (middle right), or serum/Lif from ESCs expanded in serum/Lif (right) on mouse embryonic fibroblast cells. Scale bars represent 200 μm. (D) Yield of Pdgfrα-h2b-gfp+ EBs and the number of GFP+ cells per EB in B27N2 or serum media supplemented +/− Lif and +/− RA. ANOVA with Bonferroni post-hoc test, *** p < 0.001, ** p < 0.01. (E, F, G, H, I, J) Dose-response curves showing the effect of different soluble pathway modulators after 96 h in culture on the yield of Pdgfrα-h2b-gfp+ EBs (blue), number of PrE cells per EB (red) in median focus plane (10x objective) and EB projection area (as a proxy for EB size, green). All values were normalized to H2O/DMSO controls. Average and standard deviation values were obtained from n = 3 or 4 wells with every well containing approximately 400 EBs. (K) Schematic for chemically-induced differentiation of EBs towards PrE. (L) (left) Yields for PrE differentiation (left axis) and size of EBs (right axis) using the induction cocktail. Low [c] indicates lower concentrations of 1 mM cAMP and 3 μM Chir. PrE inductions in Lif and RA/Lif media are shown for comparison. (right) Representative fluorescent images of annotated conditions. Scale bars in images and inserts represent 200 μm and 40 μm, respectively. In C, D and L bars and error bars are means and standard deviations, respectively, obtained from n = 4 wells, with each well containing ~400 EBs. *** p < 0.001, ANOVA with Tukey’s multiple comparison test. Images in panels C,D and L are taken after 96 h of culture.

### A three-dimensional screen reveals signalling pathways regulating Pdgfrα expression

Several signalling pathways influence the formation of the PrE lineage including Stat^22^, Retinoic acid^61^, Fgf^5,20,62^, Wnt^63,64^, and Tgf^65^. In the *conceptus*, these pathways are likely to act synergistically but their actions and interactions remain difficult to investigate. We thus tested these developmental pathways in cultured ESCs. While Lif (10 ng/mL) increased the yield of Pdgfrα+ EBs in serum cultures (30% yield, 3.6-fold increase, Figure 1D), the addition of Retinoic acid (RA; 10 nM) further improved the process (91% yield, 3-fold increase, Figure 1D) as well as increased the number of Pdgfrα+ clusters per EB (5.5-fold, Figure 1D, clusters are defined as Pdgfrα+ cells found within the equatorial plane of EBs, and reflecting loci of EBs differentiation into Pdgfrα+ cells, see methods and **Figure S2**). In contrast, the effect of these two molecules appeared restricted in serum-free B27N2/Lif medium (16% yield). We concluded that Lif and RA support but are not sufficient to form Pdgfrα+ cells.

We then created a small library including the known activators and inhibitors of signalling pathways active in the blastocyst (**Table 1**). We first tested them individually in serum-containing medium, and measured the percentage of Pdgfrα+ EBs (yield) and the number of Pdgfrα+ clusters per EB. The broad tyrosine phosphatase inhibitor Sodium Orthovanadate (40 μM) supported a 1.5-fold increase in yield and a 1.3-fold increase in the number of cells (Figure 1J). Fgf4 (100 ng/mL) and the Wnt activator CHIR99021 (6 μM) increased both the yield (44% and 81%, respectively) and the number of cells (both 1.6-folds; Figure 1E, 1J). Accordingly, inhibiting Wnt secretion (IWP2), Wnt processing (XAV), and MAPK (PD032) did not increase the yields (Figure 1E). We concluded that, similar to events occurring in the blastocyst^5^, the Fgf and Wnt pathways regulate both the differentiation and the expansion of Pdgfrα+ cells.

In contrast, the activation of the Tgfβ pathway by Nodal, Activin-A, BMP4 or Tgf-β1 reduced either the yield or the number of Pdgfrα+ cells. Consistent with these observations, the TGF-β receptor inhibitor SB43 and the Alk1/2 inhibitor ML347 (Bmp signalling) enhanced the formation of Pdgfrα+ cells (Figure 1I). The BMP pathway inhibitor LDN193189 prevented proliferation of ESCs (Figure 1I).We concluded that the activation of the Wnt and Fgf pathways and the inhibition of the Tgfβ pathway act on the generation of Pdgfrα+ cells.

### A three-dimensional screen reveals GPCR ligands inducing Pdgfrα expression

Next, to complement the action of the developmental pathways, we investigated the potency of GPCR ligands and ran a compound screen of 264 GPCR ligands. DL-adrenaline, a β-adrenergic agonist acting upstream of the cAMP/PKA pathway, strongly increased the yield of Pdgfrα+ EBs (206%) without affecting the overall size of the EB or the number of clones (Figure 1E). Consistent with the role of GPCR ligands in enhancing the activity of signalling pathways^66^, we concluded that DL-adrenaline potentiates ESCs for Pdgfrα expression independent of proliferation. Accordingly, the cell-permeable analogue 8Br-cAMP (3200 μM) also increased the yield of PrE EBs by 91% as compared to serum/Lif alone, Figure 1E)^55^ without affecting the size of EBs.

Altogether, we concluded that Fgf4, Wnt, Lif, RA, DL-Adrenaline, and cAMP individually increase the potential for ESCs to express Pdgfrα.

### A combinatorial screen delineates a chemically-defined medium inducing Pdgfrα expression

Because signalling molecules act in concert to ensure development, we ran combinatorials of molecules, this time in serum-free medium (B27N2 medium, **Figure S3**). Using a factorial design screening approach^67^, we tested combinations of 8Br-cAMP, DL-Adrenaline, Lif, Fgf4, Sodium Orthovanadate, Chir, ML347, SB43, RA, and Activin-A at effective concentration ranges. Specific combinations preserved EB viability (measured by EB-size), EB integrity (measured by EB circularity) and induced PrE-like differentiation (Pdgfrα expression, **Figure S4**). Among all 21 combinations, a medium containing 8Br-cAMP (1 mM), RA (10 nM), Fgf4 (100 ng/mL) and Chir (5 μM) led to a stark upregulation of the yield of Pdgfra+ EBs (78%, Figure 1K, L, **S4C**). Consistent with the important role of RA^61,65^, depleting this molecule from the induction medium reduced the yield significantly (Figure 1L). However, the synergy with other factors was essential for a robust and efficient induction (Figure 1L). This chemically-defined inductive medium also reduced the number of dead cells per EB, to levels similar to serum-containing medium (**Figure S5B**), and no longer required the presence of Lif neither for maintaining viability or Pdgfrα expression (**Figure S5A**).

### The embryoid bodies form a niche including both PrE- and Epi-like cells

The resulting EBs spontaneously formed an outer layer of PrE-like cells positive for the transcription factors Gata6^3,72–75^ and Sox17^76–78^ and a core of Nanog+ cells (96 h, Figure 2A). This spatial organization is consistent with the segregation of the PrE and Epi of the late blastocyst^5^. We then characterized the cells *via* single-cell transcriptomics (96 h). In congruence with Pdgfrα labelling, the cells showed two distinctive subpopulations (Figure 2B, **Table 2**). One expressed the PrE genes *Gata6, Gata4, Pdgfrα, Sox7 and Sox17*, while the other one expressed the Epi genes *Nanog*, *Sox2* and *Oct4* (Figure 2C). Importantly, Epi-specific Fgf4 and PrE-specific Fgfr2 were mutually expressed, which resembled the native circuitry reinforcing PrE identity in blastocysts^10,16,79^. SPRING Louvain clustering^68^ pinpointed 3 subpopulations of extra-embryonic cells (Figure 2D): one displaying the early parietal endoderm (PE) markers Follistatin and Vimentin, one opposing subpopulation displaying the early visceral endoderm (VE) markers Dab2 and Podocalyxin, while the third subpopulation had an intermediate profile. In accordance with the Louvain clustering, principal component (PC) analysis separated the embryonic and extra-embryonic cells along PC1 and sub-populations of extra-embryonic cells along PC2 (**Figure S6A, S6B**). The bottom 50 and top 50 differentially-expressed genes along the PC axis 2 showed on one side PE genes^69^ such as *Vim*, *Fst*, *Thbd*, *Sema6* and *Nog*, and VE genes^69^ such as *Amn*, *Cubn*, *Dab2*, *Podxl* (Pcx), *Apoe* on the opposite side (Figure 2E, **Figure 5SB**). In line with this, tSNE maps allowed clustering putative PE and VE subpopulations (**Figure S7A, Figure S7B)** that revealed distinct expression levels for known PE and VE genes (**Figure S7C**) and a number of potential markers for these subpopulations (Figure 2E, **Figure S6B, Table2**).

**Figure 2:**
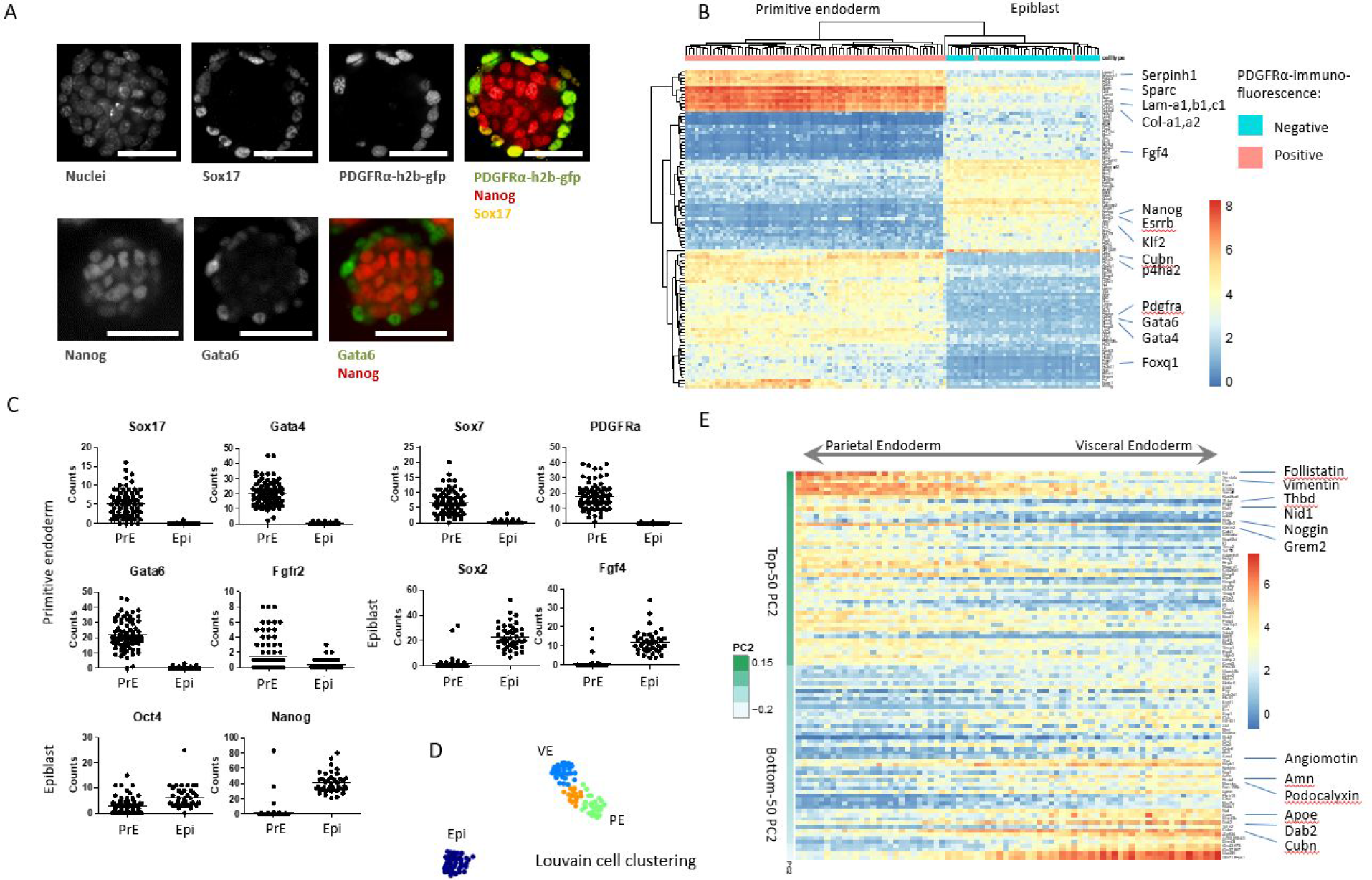
EBs form a niche including both Epi- and PrE-like cells with putative PE and VE populations. A) Immunofluorescence images of Sox17, Pdgfrα-h2b-gfp, Nanog, Gata6, and nuclei of PrE-induced EB after 96 h of culture. Scale bar represents 100 μm. (B) Hierarchical clustering of top 100 differentially expressed genes (single-cell RNA sequencing data) between Pdgfrα+ labelled cells (PrE) and Pdgfrα-cells (Epi). (C) Transcription factor RNA transcript levels for Pdgfrα+ labelled cells (PrE) and Pdgfrα-cells (Epi). (D) SPRING Louvain clustering^68^ delineates four putative subpopulations; dark blue: E4.5 Epi, light blue: early VE, orange: subpopulation of intermediate VE/PE marker profile, early PE. (E) Single-cell expression plot of Pdgfra+ depicting the top 50 and bottom 50 genes on the PC2 axis. Highlighted genes are known PE (top) and VE (bottom) genes^69–71^

From these data, we performed gene ontology term analysis (**Table 2**). In the putative VE subpopulation, cell polarity regulators typical of an epithelium and Tgf-β pathway responses were enriched as compared to Epi, in accordance with the epithelium nature of the VE and with the previously proposed function of Tgf-β in VE expansion and in Epi sustainability^45^. We conclude that the chemically defined medium induced the co-formation of spatially organized PrE-like and Epi-like cells, the former group capable of bifurcating into both VE and PE lineages.

### The PrE/Epi-like niche spontaneously progresses into extraembryonic endoderm/Epi rosettes in growth factors-free and gel-free culture conditions

*In utero*, extraembryonic endoderm (XEn)/Epi rosettes form following implantation of the blastocyst, when the VE deposits a laminin-rich basement membrane that polarizes the Epi cells and triggers the formation of the pro-amniotic cavity (Figure 3A)^80^. Such Epi-only rosettes form in the absence of PrE cells when ESCs are encapsulated in Matrigel (that mimics the basement membrane) and cultured in serum-containing medium (that provides soluble molecules)^43,81^. We thus tested whether the free-standing PrE/Epi niche was sufficient to functionally generate post-implantation-like rosettes. Upon transfer into plain B27N2 medium without growth factors, the suspended PrE/Epi structures proliferated and formed a cavity morphologically resembling the polarized XEn/Epi tissues (Figure 3A, 3B, 3C).

**Figure 3:**
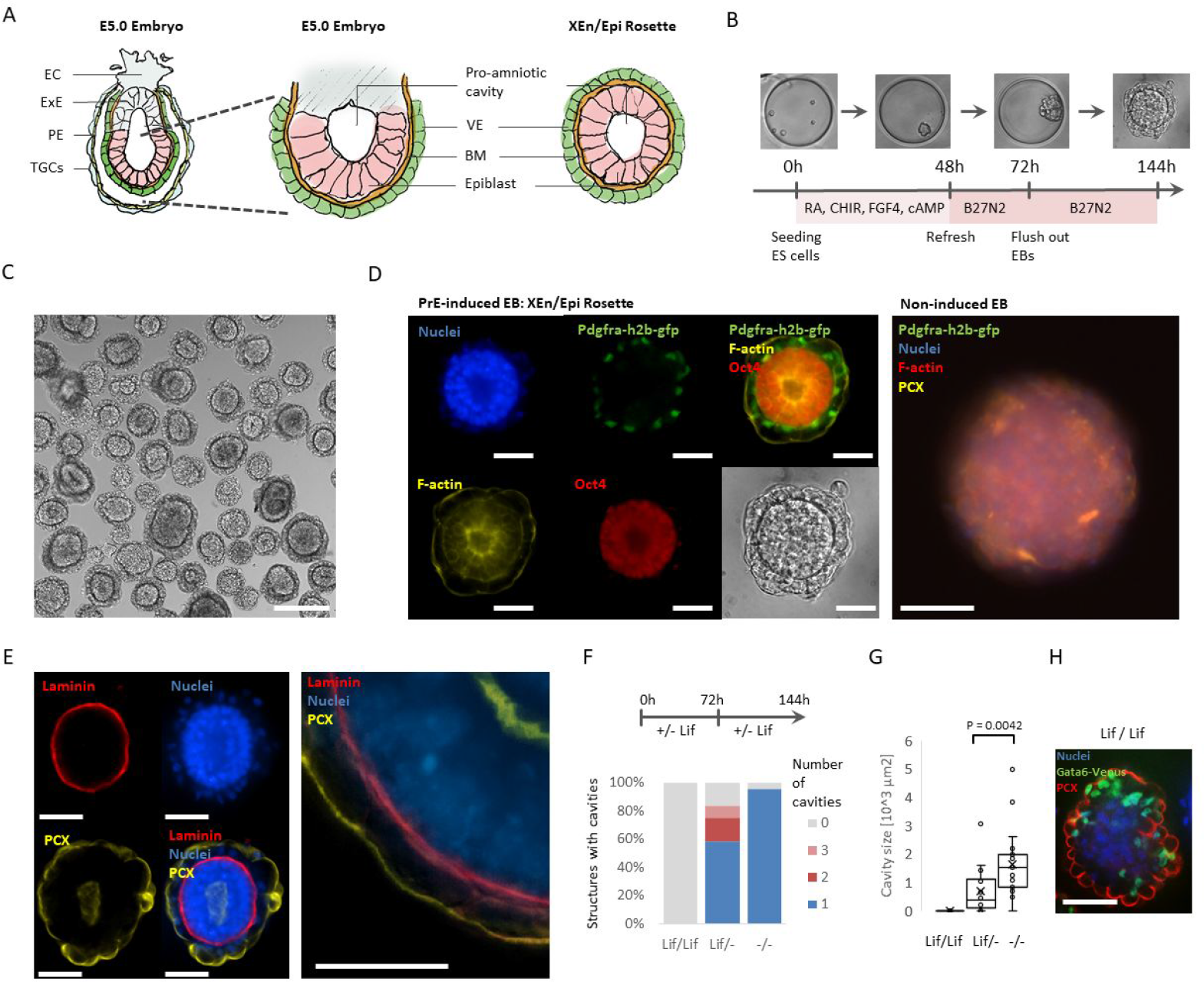
The PrE-/Epi-like niche spontaneously progresses into a post-implantation-like Endoderm/Epi rosettes in minimal culture conditions. **(A)** Schematic depicting an E5.0 conceptus (left, middle) and corresponding tissues in an XEn/Epi rosette. EC = ectoplacental cone, ExE = extraembryonic ectoderm, PE = parietal endoderm, TGCs = trophoblast giant cells, VE = visceral endoderm, BM = basement membrane. (**B)** Schematic of the culture protocol for XEn/Epi rosette formation. **C)** Brightfield image of XEn/Epi rosettes. Scale bar represents 200 μm. (**D)** Immunofluorescence and brightfield images of individual XEn/Epi rosettes images after 144 h of culture. Staining for nuclei (DNA), F-actin (pro-amniotic cavity), Pdgfrα-h2b-gfp (PrE) and Oct4 (pluripotent Epi)(left). EB cultured in the same basic conditions but without PrE-induction molecules (right). Scale bar represents 50 μm. (**E)** Immunofluorescence images depicting cell nuclei (DNA), Pcx (polarization), Laminin (basement membrane) of a XEn/Epi rosette. Scale bars represent 50 μm (**F-H)** Effect of Lif on (**F**) the percentage of structures forming a pro-amniotic cavity or multiple cavities and (**G**) the resulting cavities’ integrated surface area. Lif/Lif indicates Lif supplementation during first 3 days of PrE-induction and subsequent 3 days of XEn/Epi rosette formation, Lif/− indicates Lif supplementation only during first 3 days of PrE-induction, −/− indicates no Lif supplementation. P-value calculated according Mann-Whitney *U* test. (**H)** Immunofluorescence image of a non-cavitated and non-polarized structure resulting from Lif/Lif supplementation, labeled for nuclei, Gata6 (PrE) and Pcx (polarization). Scale bar represents 50 μm.

Interestingly, ESC alone do not proliferate in plain B27N2 medium. This suggested that the PrE- and Epi-like tissues mutually supported their proliferation. Similar to post-implantation embryos, the Epi-like cells expressed Oct4, accumulated F-actin and Pcx at the apical side to form a cavity (Figure 3D, E), while the PrE-like cells produced a laminin basement membrane and polarized as well (Pcx, Figure 3E). Over time, the cavities increased in size (**Figure S8A**). The process was both efficient (94%) and reproducible (**Figure S8B, S6C**). We concluded that the formation of the PrE/Epiblast niche was sufficient to support the spontaneous, mutual proliferation and organization of a post-implantation-like structure including a pro-amniotic cavity.

### Lif signalling inhibits the formation of the pro-amniotic-like cavity

Lif has been shown to prevent the formation of the amniotic cavity in Matrigel-embedded / serum-cultured conditions^46^. Similarly, the presence of Lif during the first 3 or for the entire 6 days of *in vitro* development respectively reduced and fully inhibited the formation of the pro-amniotic cavity, as seen by the absence of Pcx within the Epi and non-apically located Pcx in the XEn (Figure 3F-H). Also, the inhibition of apoptosis using Z-vad-fmk did not impair the hollowing of the pro-amniotic cavity (**Figure S8B)**, as previously observed^43,81^.

Altogether, we concluded that, in chemically-defined conditions, a restricted number of signalling pathways (Wnt, Fgf, RA, cAMP) are sufficient to induce the co-formation of PrE-like and Epi-like cells supporting the spontaneous organization of a structure resembling the post-implantation embryonic stage.

### The chemically-defined medium enhances Primitive Endoderm formation in blastoids

Next, we tested the capacity to modulate the induction of PrE-like cells in blastoids by exposing ESCs including a fluorescent reporter for Gata6 (ESCs^Gata6-h2b-venus/+) 82^ to the inductive molecules. For this, we primed the cells during the aggregation phase (0 – 24 hours, Figure 4A), prior to adding the trophoblast stem cells (TSCs)^56^. PrE-induction tempered the efficiency of blastoid formation (from 49% to 36%, Figure 4B) by reducing the efficiency of TSCs to engulf the EBs (from 39% to 30% of non-engulfed structures, Figure 4B). Nevertheless, the induction increased the overall percentage of blastoids including Gata6+ and Nanog+ cells to 78%, compared to 22% under standard conditions (Figure 4D, E). Concomitantly, the number of Gata6+ cells increased (p-value = 0.00079, Figure 4E, F). Notably, the total number of inner cells was also higher upon PrE induction (Figure 4G). In accordance with our observations in rosettes and with a previous study^83^, these data confirm that a synergy between PrE and Epi cells regulates the total number of inner cells. The ratio of Gata6+/ Nanog+ cell numbers in PrE-induced blastoids was comparable to the one in blastocysts (0.83 vs. 0.9 in 120 cells-stage blastocyst ((Saiz et al. 2016), Figure 4H).

**Figure 4:**
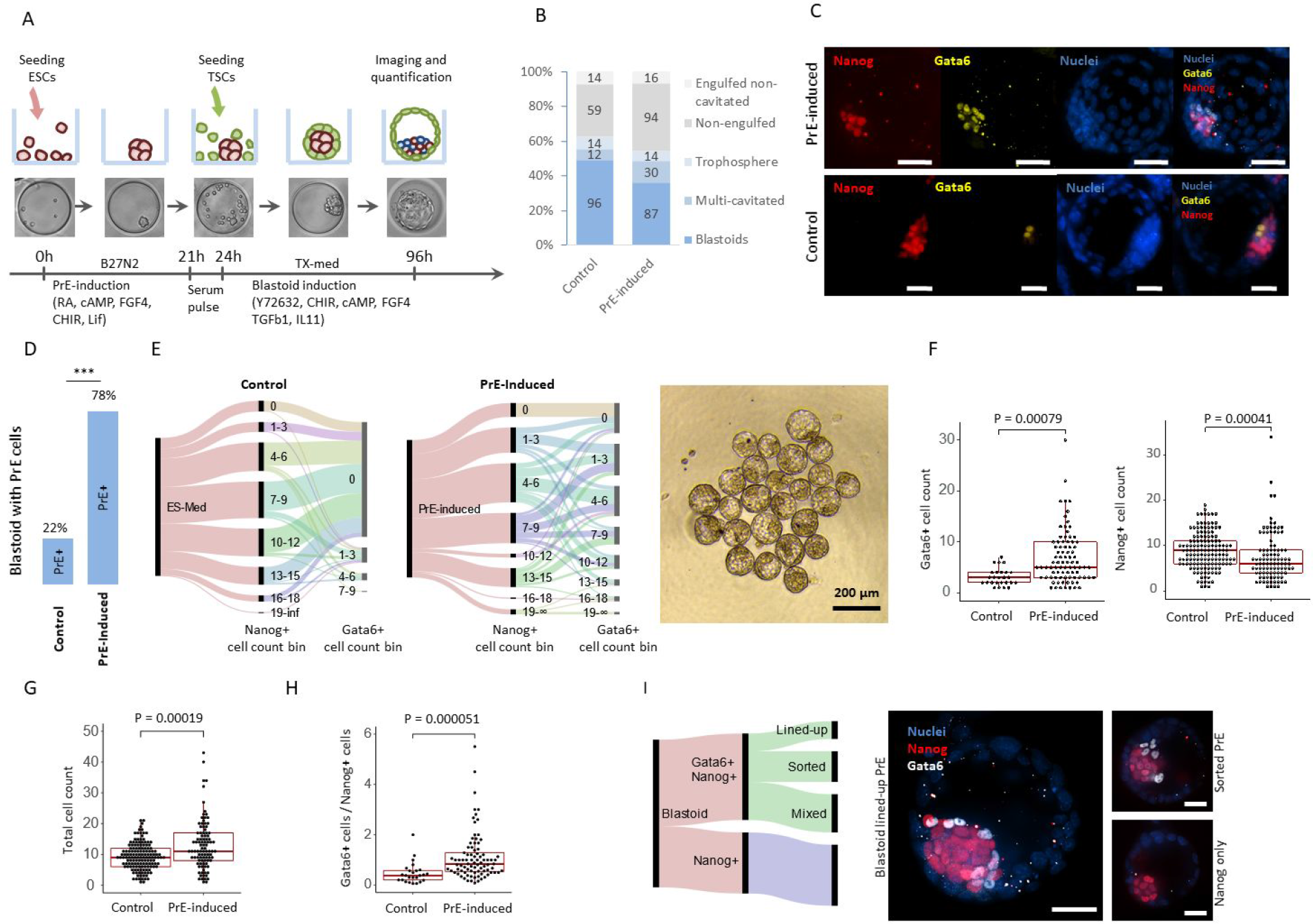
The PrE/Epi priming of ESCs induces the formation of the niche in blastoids. **(A)** Schematic of experimental design and stage-specific bright field images of PrE-induced blastoid formation. (**B)** Percentages of the different structures found in microwell arrays in control (n=195) and PrE-induced (n=241) conditions. (**C)** Representative immunofluorescence images of PrE-induced and control blastoids stained for Nanog (red) and Gata6 (yellow). DAPI staining (blue) shows cell nuclei. Scale bar represents 50 μm. **D)** Percentage of blastoids including Gata6+/Nanog+ cells formed under control of PrE-induced conditions (left).*** p < 0.001, Fisher’s exact test. (**E)** Alluvial diagram displaying cell count of Nanog+ and Gata6+ dichotomy for control and PrE-induced blastoids (left). Brightfield image of a selection of PrE-induced blastoids (right). (**F)** Gata6+ and Nanog+ cell counts compared between control and PrE-induced blastoids that contain both Nanog+ and Gata6+ cells. (**G)** Total number of inner cells within blastoids (sum of Gata6+ and Nanog+ cells). (**H**) Ratio of Gata6+ /Nanog+ cells per blastoid containing both Gata6+ and Nanog+ cells. (**I)** Alluvial diagram displaying contributions of resulting phenotypes following PrE-induced blastoid formation. Representative immunofluorescence images of blastoids with; a lined-up Gata6+ cell epithelium on top of Nanog+ cell cluster (left), sorted PrE but not lined-up (top right) and Nanog+ cells only (bottom right). Images acquired by stacking 3 to 5 spinning disk confocal slides using a 40× objective. Scale bar represents 50 μm. In F, G and H the P-values were determined by the Mann-Whitney *U* test.

Next, we examined the spatial organization of PrE-induced blastoids. When blastocysts progressed, the PrE cells sorted out from the Epi cells to line the blastocoel cavity. We observed that 21% of the PrE-induced blastoids including PrE-like cells showed a layer of Gata6+ cells lined-up along the cavity of the blastoid (Figure 4I)^10,84^. Among the other blastoids, 35% comprised sorted but not lined-up Gata6+ cells, while 44% had a salt and pepper phenotype of Gata6+ and Nanog+ cells (Figure 4I) reminiscent of an earlier blastocyst stage^7,32,85^

### The induction of Primitive Endoderm supports the in vitro formation of post-implantation-like structures from blastoids

We then tested whether the PrE/Epi-like tissue within blastoids could support the development of structures reflecting aspects of the post-implantation *conceptus*, such as the 3-dimensional organization of the epiblast. We plated PrE-induced blastoids containing PrE cells (>2 Gata6+ or Pdgfrα+ cells) and non-induced blastoids *in vitro*^*86,87*^ The presence of PrE cells (Gata6+) enhanced the potential to expand the PrE (96 h, Gata6+, 53% vs. 10%, Figure 5A). Similar to our previous observation^56^, PrE-induced blastoids with Gata6+ cells and non-induced blastoids maintained survival of Epi cells (96 h, 98% vs 100%, Figure 5A).

**Figure 5:**
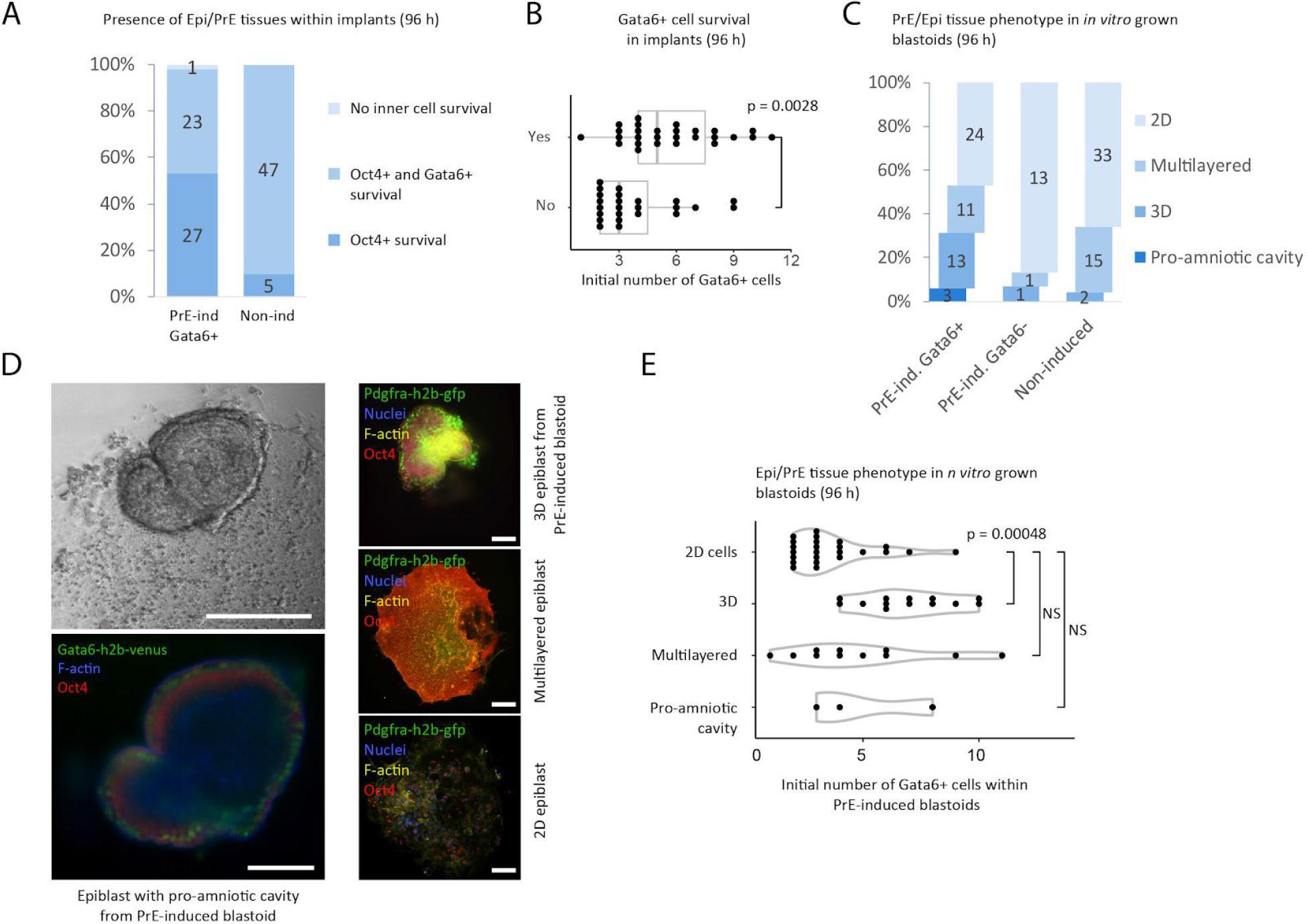
The induction of the PrE-/Epi-like niche in blastoids supports the formation of post-implantation-like tissues. (A) Survival of Epi (Oct4+) and PrE (Gata6+) tissues within *in vitro* grown PrE-induced blastoids with Gata6+ cells (96 h). Total number of structures are displayed within bars. **(B**) The presence of PrE tissue (Gata6+) within *in vitro* grown PrE-induced blastoids (yes/no, at 96 h) as a function of the numbers of Gata6+ cells within the initial blastoids. P-value by Mann-Whitney *U* test. **(C)** Percentage of different tissue phenotypes from PrE-induced blastoids including or not Gata6+ cells as compared to non-induced blastoids. **(D) (left)** Brightfield and immunofluorescence images of an *in vitro* grown blastoid with Oct4+ Epi (red) and Gata6+ PrE (green) cell surrounding a pro-amniotic cavity and growing on top of a TSC monolayer (96 h). **(right)** Representative images of XEn/Epi tissues phenotypes with Oct4+ Epi (red), Pdgfra+ PrE (green) and overall F-actin (yellow) and nuclei (blue). Scale bars represent 200 μm. **(E)** Tissues phenotypes as a function of the number of Gata6+ cells within PrE-induced blastoids. Kruskal-Wallis *H* test followed by a pairwise Mann Whitney *U* test with Bonferroni correction.

We observed that the potential for the PrE cells to expand depended on the initial number of PrE cells present in blastoids (Figure 5B). In addition, the induction of PrE cells improved the formation of 3D structures (35% vs. 7%, Figure 5C) containing both Epi and PrE cells. Consistent with our findings in EBs, we concluded that PrE cells are critical for the proliferation and organization of the post-implantation Epi. Clearly, the potential of the Epi to expand and initiate the formation of a 3D structure depended on the initial number of PrE cells present in blastoids (Figure 5E). We concluded that PrE-induction promoted the survival of expandable Epi- and PrE-like cells. Finally, upon culture, the PrE-induced blastoids supported the formation of rosettes including Gata6+/ Pdgfrα+ cells surrounding Epi-like cells and including a pro-amniotic-like cavity (2 out of 14 and 1 out of 9 PrE-induced blastoids in two separate experiments, Figure 5D). In sharp contrast, the non-induced blastoids lacked that potential.

Altogether, we concluded that a specific combination of signalling pathways is sufficient to rapidly and efficiently generate a PrE/Epi-like niche within blastoids that supports the *in vitro* expansion and morphogenesis of a post-implantation-like tissues.

## Discussion

High-content screening of a large number of embryoid bodies on a microwell array in chemically-defined culture conditions allows for robust statistics necessary to delineate the effect of signalling pathways. Here, we observed that the combination of cAMP, RA, Wnt and FGF4 signals were sufficient to rapidly and efficiently drive the co-formation of PrE-like cells and Epi-like cells in gel-free and serum-free cultures. Upon transfer in plain medium, these two cell types are capable of further growth and autonomous organization into a structure resembling post-implantation embryonic and extraembryonic tissues.

Consistent with the idea that the blastocyst is a self-regulating system^83^, the induction of PrE-like cells in blastoids shows a synergy between PrE and Epi cells regulating the total cell number, which sustains the expansion and morphogenetic capability during the post-implantation stage. Accordingly, the insufficient formation of PrE and incomplete lining of the PrE epithelium between the Epi and blastocoel has been described to halt Epi expansion in blastocysts^59,81^. Similarly, inappropriate specification of the extraembryonic VE has been shown to result in disorganized ectoderm and stagnated development ^88^. Overall, these results argue for the importance of the PrE tissue in nurturing the Epi for survival and expansion, and for a synergistic development of the embryonic and extraembryonic tissues. The mechanisms by which the PrE accomplishes this remain to be determined.

Altogether, this study establishes stem cell-based models of the embryo amenable to high-throughput drug and genetic screens and alleviating the burden on the use of animals^57,89^. We propose that these models are foundations for basic and biomedical discoveries to elucidate the critical and currently unknown processes of embryogenesis.

## Methods and Materials

No statistical methods were used to predetermine the sample size.

### Microfabrication

Elastomeric stamps for imprinting the agarose microwell arrays were fabricated using PDMS Sylgard 184 kit. Microwell arrays were moulded as described previously^58^ using a 2.2% w/v solution of Ultrapure agarose (Thermo Fisher Scientific 11560166). In each well of a 96-well-plate 430 microwells, with a diameter of 200 μm, were moulded. Each well contains a calculated liquid volume of 250 μL split between 225 μL medium and 25 μL hydrogel buffer.

### Stem cell culture

The following lines were used for experiments: Pdgfrα h2b-GFP/+, H2B-RFP V6.5 sub-clone, Gata6 H2B-Venus/+;ColA1 TetO-Gata4-mCherry/+;R26 M2rtTA/+ ES cells. Gata6 H2B-Venus cell line was a kind gift of C. Schröters laboratory. The V6.5 cell line has a C57BL/6 × 129/Sv background and was obtained from the laboratory of R. Jaenisch. The Pdgfrα H2B-GFP/+ cell line has an ICR background and was derived from A.-K. Hadjantonakis’ laboratory. Standard ES cell expansion was done in 2i/Lif conditions comprising B27N2 (Gibco) medium with leukaemia inhibitory factor (Lif, Merck Millipore ESG1106, 10 ng/mL), PD0325901 (1 μM, AxonMed 1408) and CHIR99021 (3 μM, AxonMed 1386) as developed previously (Nichols et al. 2009), supplemented with 50 μM 2-mercaptoethanol (Gibco 11528926).

ESCs were seeded and expanded as 25.000 per cm^2^ on 0.1% w/v gelatin-coated tissue-culture treated polystyrene dishes (Nunc). ESCs were expanded for minimally 2 passages before aggregating into EBs within the agarose microwells. For cell banking and serum/Lif experiments ESC were expanded on a monolayer of mouse embryonic fibroblasts (mEF) in DMEM containing sodium pyruvate and Glutamax (Thermo Fisher, 10569010) supplemented with 10 mM Non-essential amino acids (Gibco, 12084947), 10 mM HEPES (Gibco, 15630056), 10 ng/mL Lif and 50 μM 2-mercaptoethanol

TSCs were seeded and expanded as 25.000 per cm^2^ on 3% Matrigel-coated dishes in chemically defined TX-medium as developed previously (Kubaczka et al. 2014) or on TCPS with a Laminin-512 coating (Biolamina LN521-02, 5 μg/mL overnight incubation at 4 degree C) in TX-medium supplemented with Activin-A (Bio-techne 338-AC-010), 50 ng/mL IL11 (Peprotech, 220-11), 200 μM 8Br-cAMP (Biolog, B007-50E), 25 ng/mL BMP7 (RnD systems, 5666-BP-010), 5 nM LPA (Tocris 3854), 2 ng/mL TGFβ1 (Peprotech, 100-21-B), 25 ng/mL Fgf4 (RnD systems, 5846-F4-025), 1 μg/mL heparin (Sigma), 100 μM 2-mercaptoethanol (Gibco 11528926).

### EB, PrE/Epi and XEn/Epi formation

EBs were formed by seeding an average of 7 ESC per microwell in either serum medium, Lif/serum medium or B27N2-based media, all supplemented with 50 μM of 2-mercaptoethanol. Serum medium consists of high-glucose DMEM containing sodium pyruvate and Glutamax (Thermo Fisher, 10569010) and supplemented with 10 mM Non-essential amino acids, 10 mM HEPES and penicillin/streptomycin. Lif was supplemented in a standard working concentration of 10 ng/mL. PrE-induction medium consists of advanced B27N2 medium supplemented with 3 μM CHIR99021, 50 ng/mL Fgf4, 10 nM RA, 1 mM 8Br-cAMP and 50 μM 2-mercaptoethanol.To induce XEn/Epi Rosettes from PrE-induced EBs, 14 ESCs were seeded per microwell. Medium was refreshed after 48 h of culture with advanced B27N2 supplemented with 50 μM 2-mercaptoethanol. After 72 h EBs were flushed out and transferred into 6-well plates with 2 mL advanced B27N2 medium supplemented with 50 μM 2-mercaptoethanol.

### Blastoid formation

Blastoids were formed as described previously^56,90^. For control blastoids, an average of 7 ESC was seeded per microwell in serum/Lif medium (10 ng/mL Lif, control blastoids) with 50 μM 2-mercaptoethanol. For PrE-induced blastoids an average of 7 ESC was seeded per microwell in either 1) B27N2 medium with PrE-induction compounds and 10 ng/mL Lif for 21 hours incubation followed by serum/Lif for 3 hours, or 2) Serum/Lif medium with PrE-induction compounds. After 24 hours of ESC aggregation an average of 17 TSC were added per microwell in TX-medium with non-essential amino acids and the blastoid culture components; 20 μM Y27632 (AxonMed 1683), 5 μM CHIR99021 (AxonMed 1386), 1 mM 8Br-cAMP, 25 nG/mL FGF4, 15 nG/mL TGFβ1, 30 nG/mL IL11, 1 μg/mL heparin, 100 μM 2-mercaptoethanol. After 24 hours an additional 1 mM of 8Br-cAMP was added to the blastoid culture medium.

### *In vitro* post-implantation assay

Blastoids cultured for 96 h (from seeding ESC) were selected on their morphology (cystic, roundness, presence of inner cell mass) and transferred from microwells onto tissue-culture glass or polystyrene plastic in IVC1 medium using mouth pipetting. IVC1 medium consisted of Advanced DMEM/F12 medium (Fisher Scientific, 11540446) with non-essential amino acids and sodium pyruvate, 10% ESC-selected fetal bovine serum, 1/100 Glutamax (Fisher Scientific, 35050061), penicillin/streptomycin, 1/100 ITS-X (Fisher Scientific, 10524233), 8 nM β-estradiol (Sigma, E8875), 200 ng/mL progesterone (Sigma, P8783), 50 μM 2-mercaptoethanol (Gibco 11528926). Structures were fixated after 96 hours of culture using a fresh solution in PBS of 2% paraformaldehyde and 0.1% glutaraldehyde.

### Image-based analysis

Fluorescence images were acquired on a widefield Nikon Eclipse Ti microscope using a 10x objective. Analysis of EBs was performed using a custom-made pipeline in CellProfiler2.0 (Broad Institute)^91^. The number of EBs and cells positive for Pdgfrα-h2b-gfp were determined by thresholding on intensity. EBs were considered positive for Pdgfrα when one or more cells were positive for Pdgfrα-h2b-gfp. The number of Pdgfrα+ cells was identified from widefield images acquired within the equatorial plane of EBs, and therefore reflect a proxy for the total number of Pdgfrα+ cells per EB..

### Soluble factor screening

Serial dilutions of compounds were made in appropriate solvents (DMSO or H2O) and corresponding carrier controls were included in the assays. Serial dilutions were made for single soluble factor titrations; FGF4, Sodium (Ortho)Vanadate (Sigma Aldrich S6508), PD0325901, PD98059 (Sigma P215), Activin-A, TGF-β1, A83-01 (Tocris 2939), Nodal (R&D systems 1315-ND-025), SB431542 (Tocris 1614), Retinoic Acid (Sigma R2625), DL-Epinephrine HCl (Sigma E4642), 8Br-cAMP, SC144 (R&D systems 4963 /10), IL6 (Peprotech 216-16), IL11, Lif, BMP4 (Peprotech 315-27), BMP7, LDN 193189 (Tocris 6053), ML347 (Selleckchem S7148), Noggin (Peprotech 250-38), CHIR99021, IWP2 (Selleckchem S7085) and XAV-939 (Selleckchem S1180).

### Immunofluorescence

Blastoids and blastocysts were fixed in 4% paraformaldehyde solution in 1X PBS for 15 minutes at room temperature. VE/Epi rosettes and *in vitro* implantation cultures were fixated in 2% paraformaldehyde solution with 0.1% glutaraldehyde in 1X PBS for 15 minutes at room temperature. After fixation, samples were washed 3x in washing buffer (0.1% Triton-X with 2% BSA in 1×PBS), permeabilized in 1% Triton-X solution in 1X PBS and blocked in blocking buffer (2% BSA, 5% serum of host 2nd antibody species, 0.5% glycine, 0.1% Triton-X, 0.2% Tween-20) for 30 minutes. Samples were incubated in antibody solution (¼ of blocking buffer with ½ of 1×PBS and ¼ of 0.1% Triton-X in 1×PBS) with primary antibodies for 12 hours at 4 degree C, washed 3x 10 minutes with washing buffer followed by incubation with secondary antibodies in antibody solution for 4 hours at 4 degree C. Optionally complemented with DAPI (0.2 μg/mL) and phalloidin (Thermo Scientific, 1/100 dilution).

### Antibody list

Mouse Podocalyxin, RnD systems AF1556, 1/300 dilution

Human/mouse Oct-3/4, RnD systems AF1759, 1/150 dilution

Human Sox17, RnD systems AF1924, 1/150 dilution

Anti-Laminin antibody produced in rabbit, Sigma Aldrich L9393, 1/100 dilution

Anti-Nanog antibody (Abcam ab80892), 1/150 dilution

Anti-Gata6 (mouse/human) poly Goat (AF1700) 200uG/mL, 1/200 dilution

Donkey anti-Goat IgG (H+L) Secondary Antibody, Alexa Fluor 647 conjugate, 1/500 dilution

Donkey anti-Goat IgG (H+L) Secondary Antibody, Alexa Fluor 568 conjugate, 1/500 dilution

Goat anti-Rabbit IgG (H+L) Secondary Antibody, Alexa Fluor 647, 1/500 dilution

Goat anti-Rabbit IgG (H+L) Secondary Antibody, Alexa Fluor 568, 1/500 dilution

### Data analysis and reproducibility

Sample sizes and statistical tests for every experiment are annotated in the figure legends. Sample sizes were not predetermined using statistical methods. If not stated otherwise, all data are displayed as mean ± the standard deviation.

PrE-induced blastoid formation and in vitro implantation assays were repeated at least five times and using two ESC and two TSC lines.

Alluvial figures were created using RAW - an open source project by DensityDesign Lab and Calibro^92^ Single-cell transcriptome analysis was performed using the Seurat package for R (https://satijalab.org/seurat/). A minimal detection threshold of 5000 genes per cell was selected for cells to be included for analysis. Clustering analysis and heatmaps were made in R^93^ using the Seurat package. Bar plots were made using Microsoft Excel. SPRING Louvain clustering was performed using Kleintools^68^.Scatter plots in Figure 2 were generated using GraphPad Prism 5. Scatter plots in Figure 4, violin plots and dose-response curves were generated using R with the packages “ggplot2”, “reshape2”, “ggsignif” and “ggbeeswarm”. Statistical analysis was performed using the package “stats”.

## Data availability

The single cell transcriptomic dataset generated and analyzed during the current study have been deposited at the NCBI’s Gene Expression Omnibus repository(Edgar et al. 2002) and is accessible through GEO Series accession number GSE129655 (https://www.ncbi.nlm.nih.gov/geo/query/acc.cgi?acc=GSE129655). All other data supporting the findings of this study are available from the corresponding author upon reasonable request.

## Acknowledgements

The authors would like to thank Christian Schröter for providing the Gata6-Venus mouse ES cells, Valerie Prideaux, Jodi Garner and Janet Rossant for providing the F4 mouse TS cell lines and Anna-Katerina Hadjantonakis for providing the H2B-GFP-Pdgfrα mouse ES cells.

