## Supplementary figures for "Chemically-defined induction of a primitive endoderm and epiblast-like niche supports post-implantation progression from blastoids"

### Supplementary information

### S1

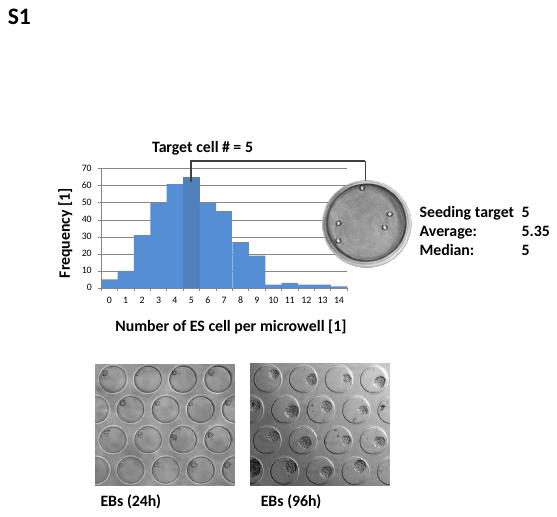

Distribution of ESC numbers within microwells with a target seeding number of 5 cells per microwell (top). Brightfield images of EBs at 24 and 96 h of culture.

### S2

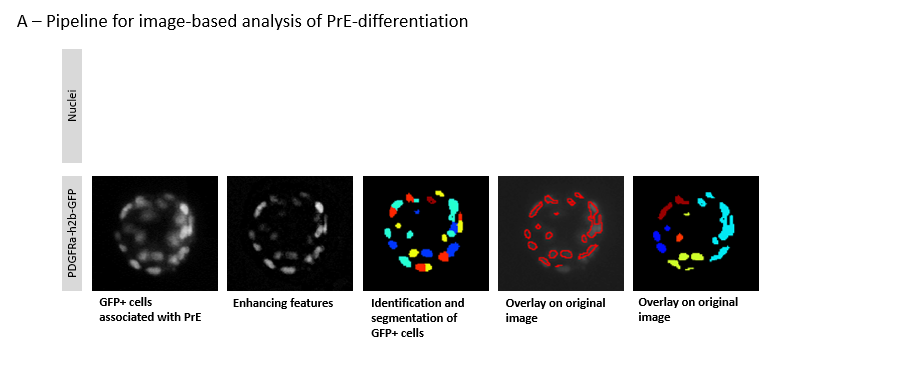

Pipeline for identification, segmentation and quantification of GFP+ cells within EBs.

### S3

Fluorescent montage images of EB cultures within hydrogel microwell screening arrays. Red color depicts the projection area of EBs identified by nuclear staining (Dapi). Green color depicts Pdgfra-h2b-gfp reporter expression.

A) B27N2 2i/lif expansion followed by serum/lif EB culture.
B) B27N2 2i/lif expansion followed by B27N2/lif EB culture.
C) Serum/lif expansion on mEF followed by B27N2/lif EB culture.
D) Serum/lif expansion on mEF followed by serum/lif EB culture.

### S4A Combinatorial screening for serum-free differentiation into primitive endoderm.

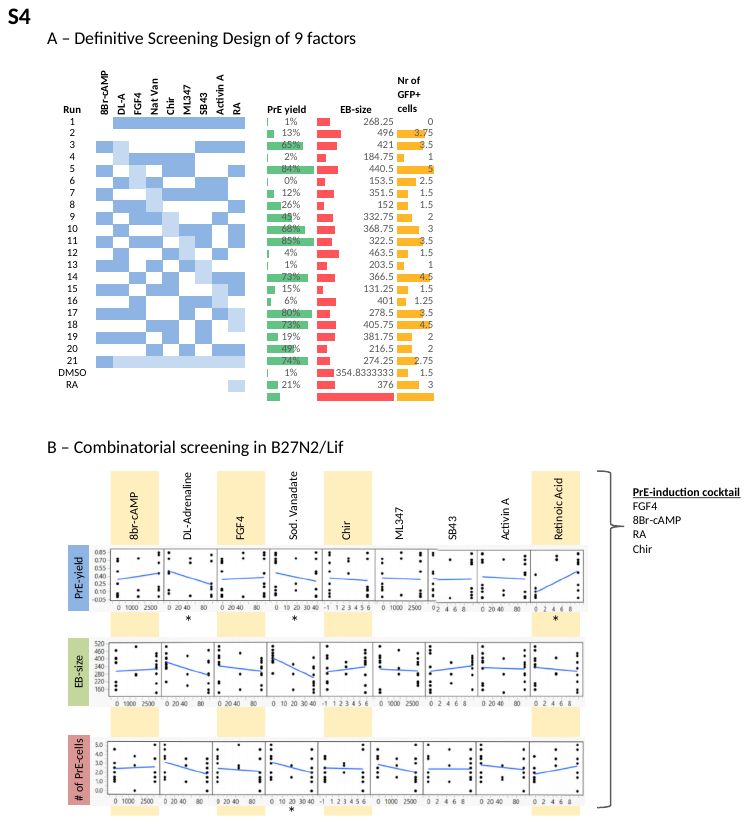

Experimental layout of Definitive Screening Design and the results of factor combinations on the yield of PrE-differentiation in EBs (PrE-yield), EB projection area (EB-size) and number of GFP+ cells per EB (Nr of GFP+ cells).

### S4B

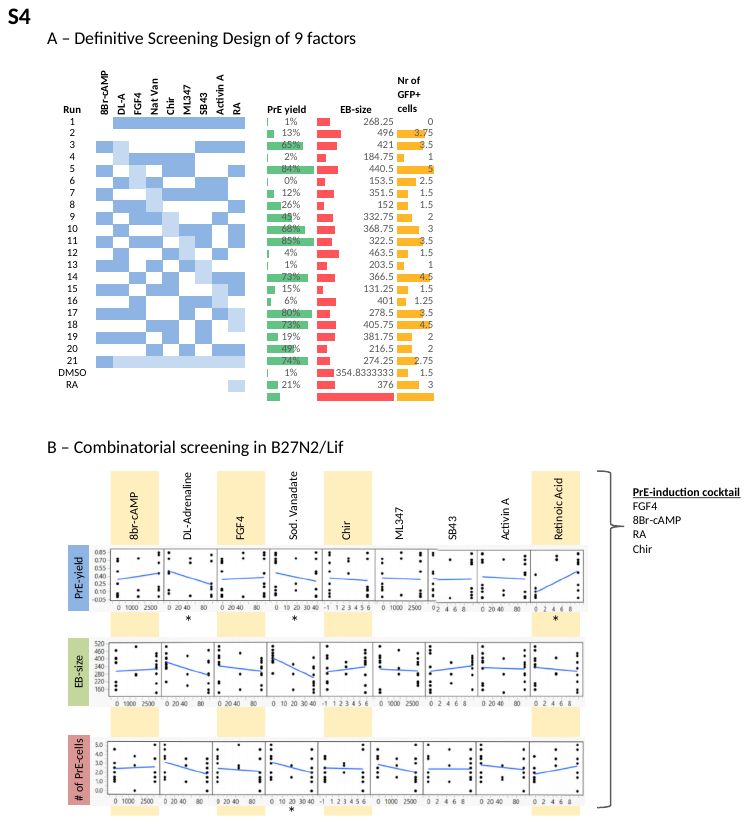

Main effect plots for the 9 factors in the combinatorial definitive screening design (DSD) assay shows a positive correlation with PrE-yield for 8Br-cAMP, FGF4 and RA, a positive correlation with EB-size for Chir, SB43 and 8Br-cAMP, and a positive correlation with the number of PrE+ cells for RA. All conditions were supplemented with Lif and β-mercaptoethanol. Asterisks indicate statistical significance. Yellow-marked compounds; FGF4, 8Br-cAMP, RA and Chir were selected for final induction cocktail.

### S4C

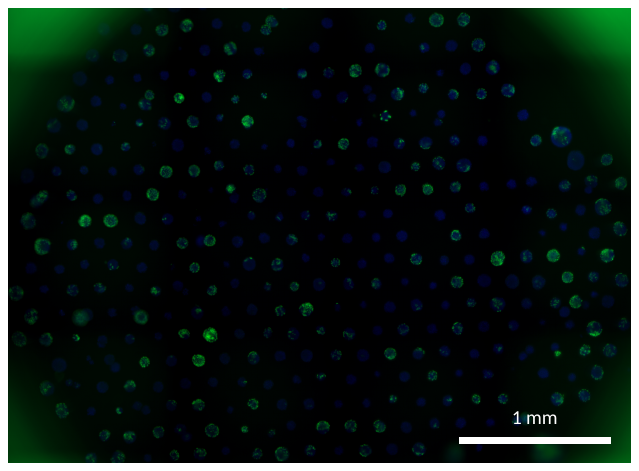

Cropped fluorescent montage image of EB cultures within a single well of a 96-wellplate that was induced for primitive endoderm differentiation using the PrE-induction cocktail (image readout at 96 hours of EB culture). Blue color indicates labelling of cell nuclei (Dapi). Green color depicts Pdgfra-h2b-gfp reporter expression.

### S5

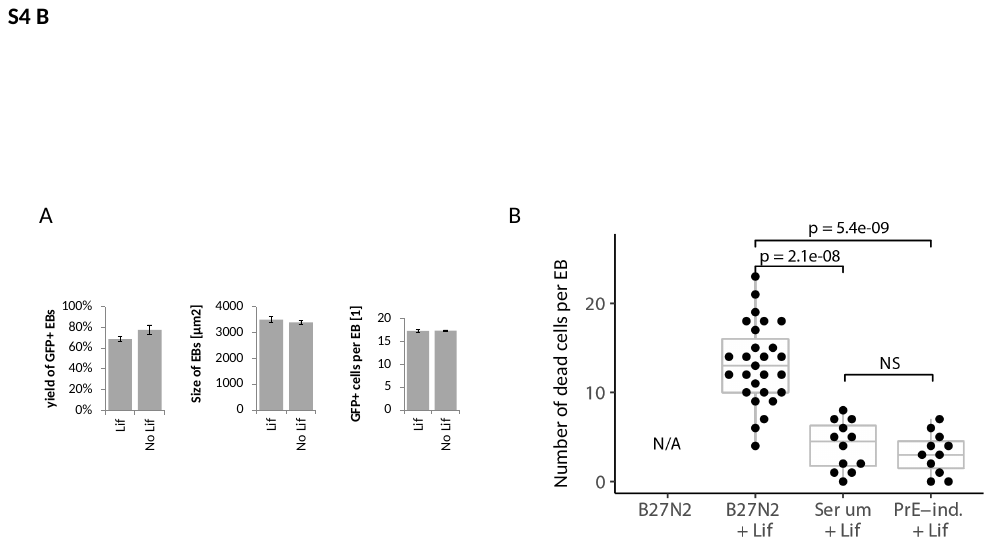

A) Yield of Pdgfrα++ EBs, size of EBs and number of Pdgfrα+ cells per EB with the PrE induction cocktail with and without Lif.
B) B) Cell viability assay; quantifying the number of dead cells (Ethidium homodimer-positive cells) per EB comparer for standard per EB after culture in the different media: serum-free B27N2 without and with Lif, serum with Lif, and the PrE-induction cocktail including Lif. EB culture Serum + Lif and serum-free B27N2 minus Lif, with Lif and with PrE-induction cocktail including Lif.

### S6A

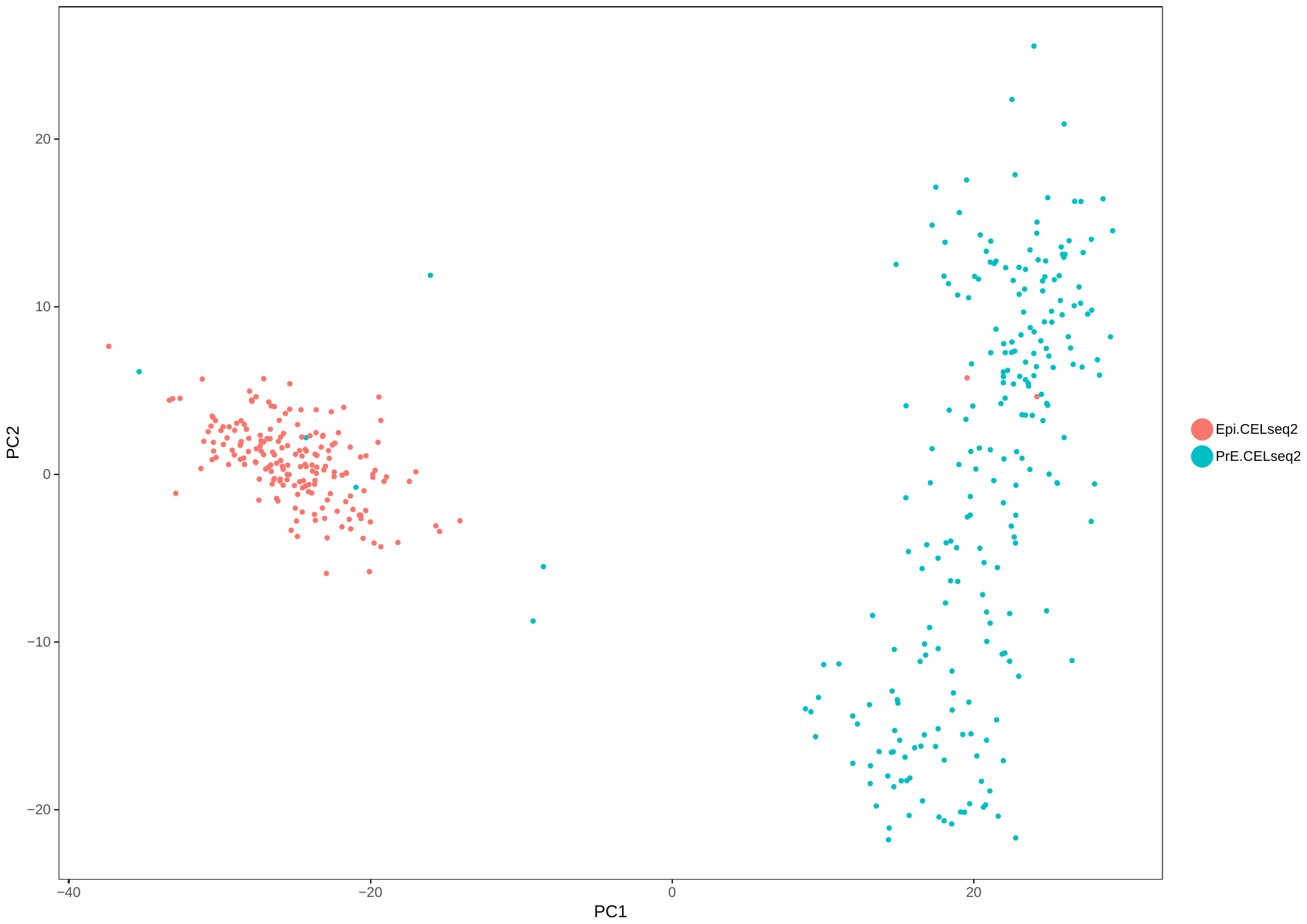

Principal component analysis on snSeq data of PrE-induced EBs.

### S6B

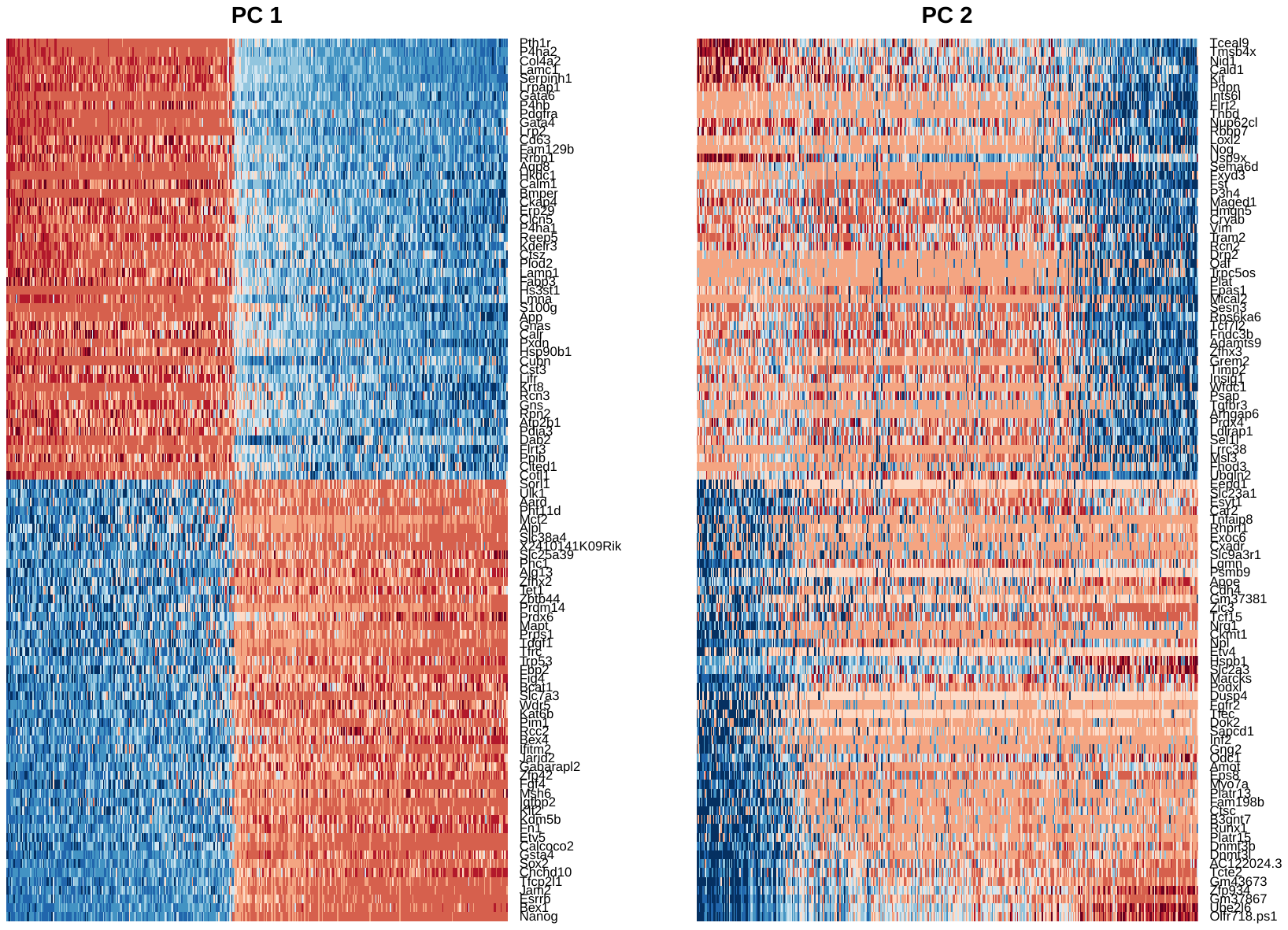

Heatmap of scaled data ordered by PC value.

### S7A

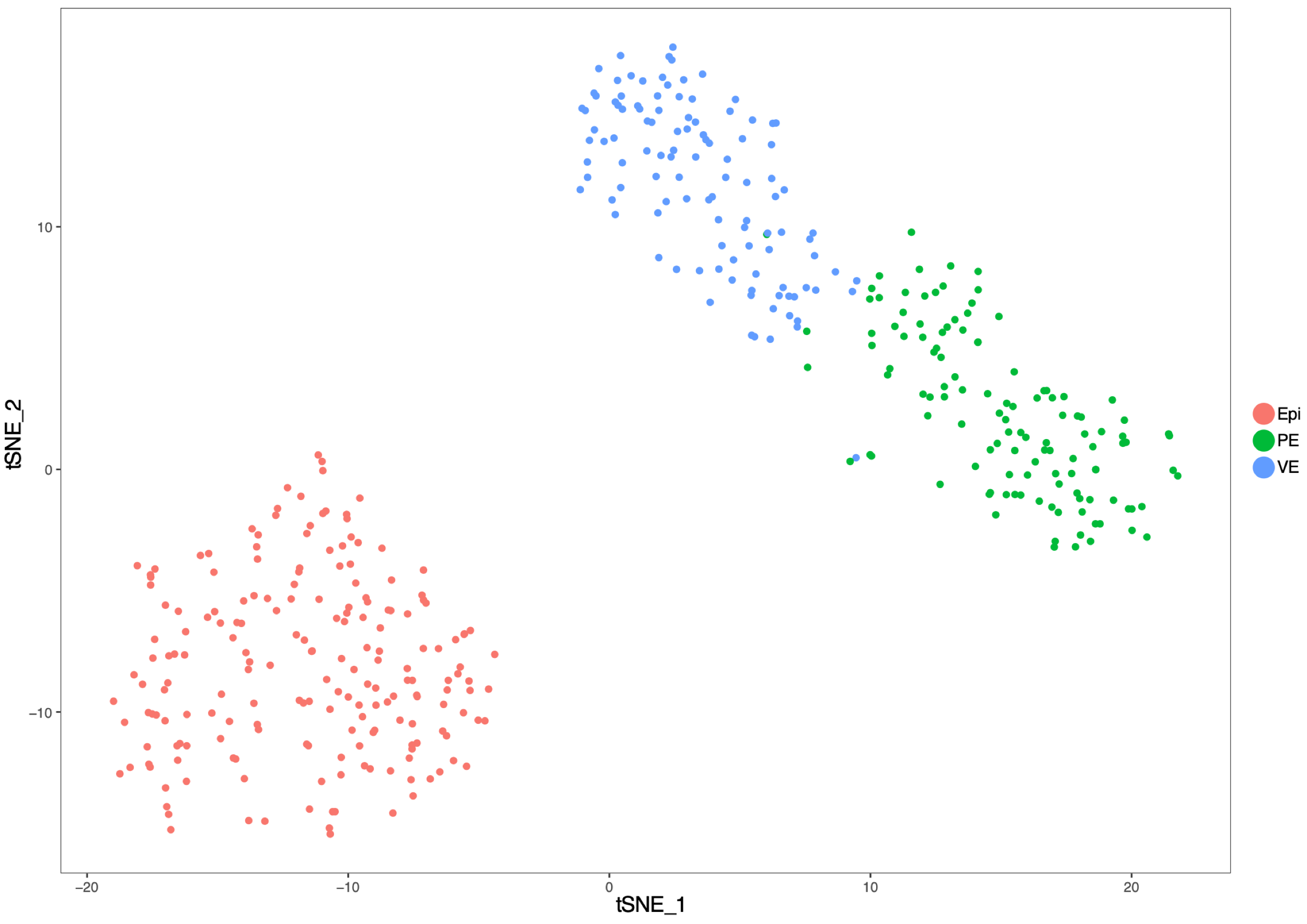

tSNE map identified three subpopulations of putative early VE, putative early PE and Epi.

### S7B

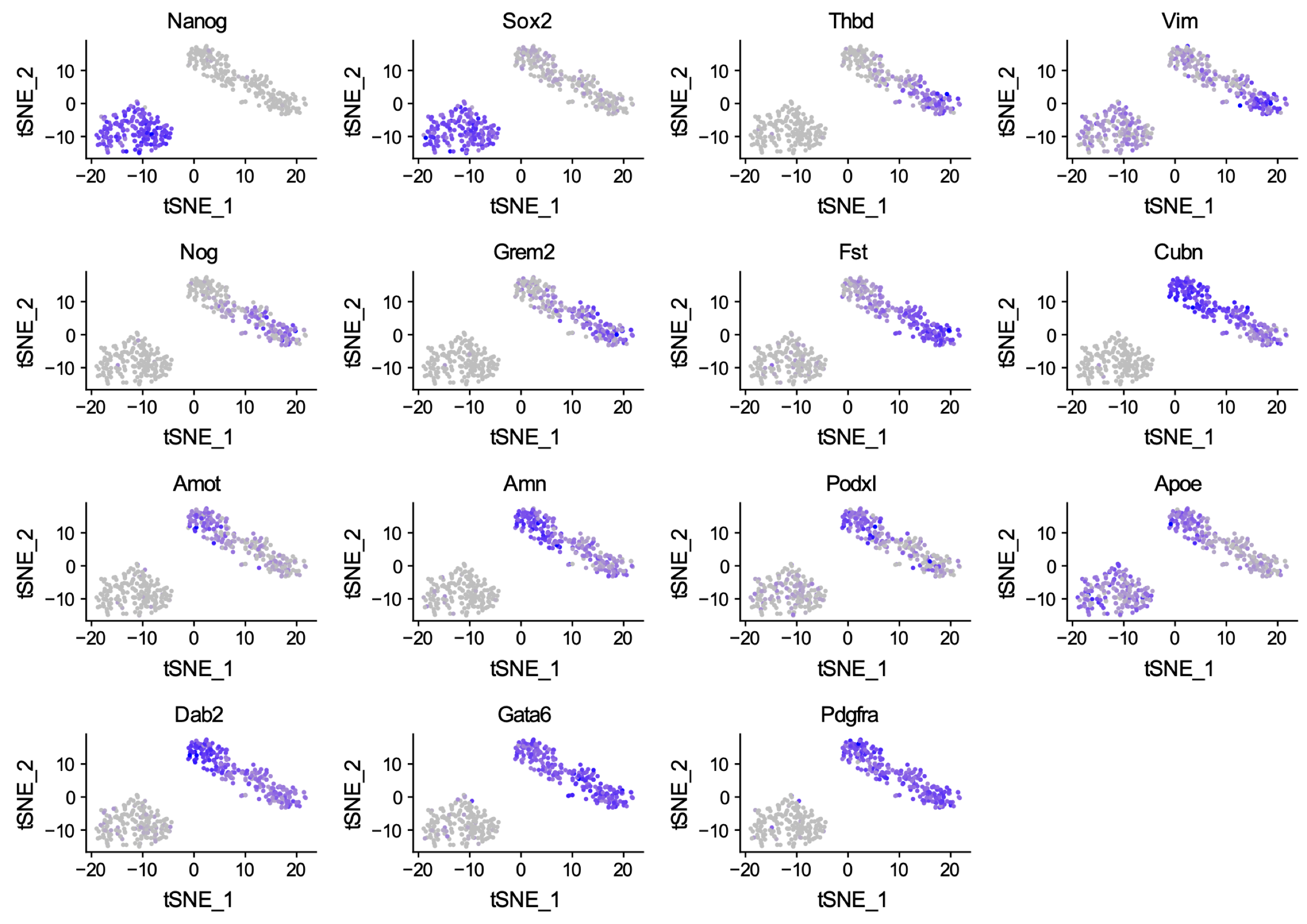

tSNE maps for VE and PE genes.

### S7C

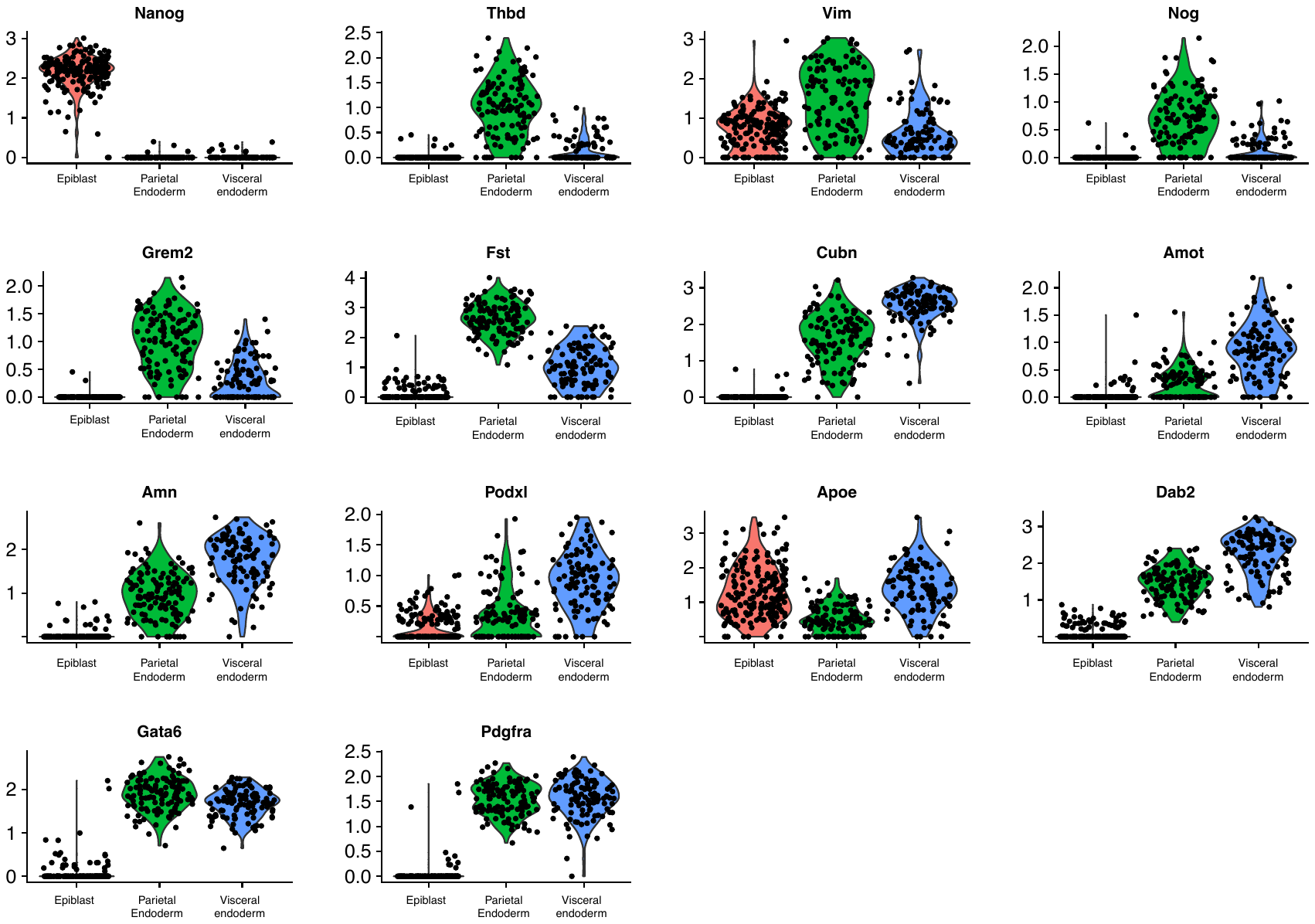

Relative expression levels for VE and PE genes within the three subpopulations (putative early PE, putative early VE and Epi)

### S8

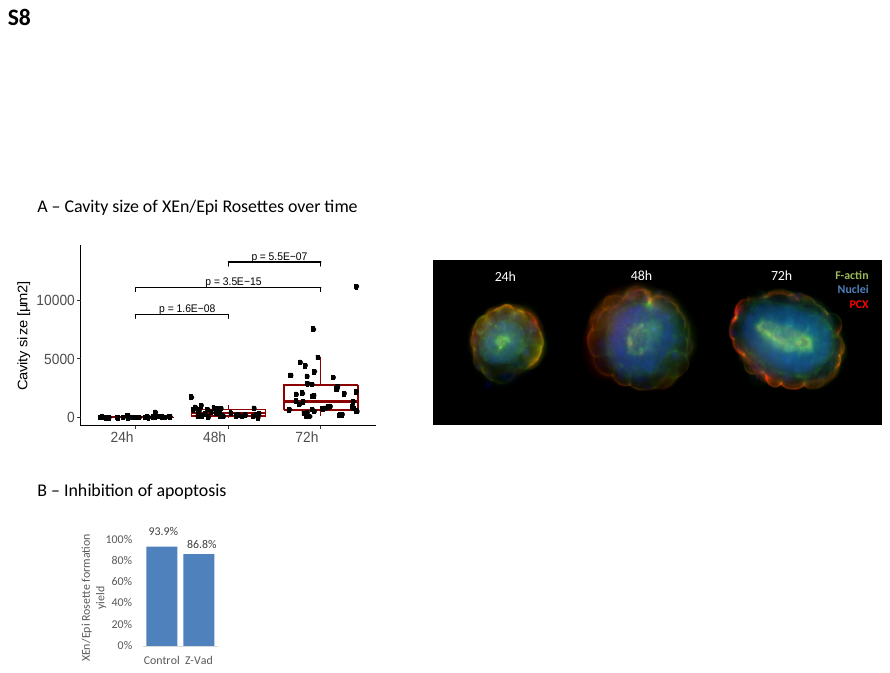

A) High-throughput formation of XEn/Epi rosettes using PrE-induction factors in B27N2 medium (average of 24 ESCs per microwell, brightness-adjusted bright-field image). B) Size of cavities in XEn/Epi Rosettes over time. B) Formation efficiency of XEn/Epi Rosettes containing pro-amniotic-like cavities with and without addition of apoptotic inhibitor Z-vad-fmk (Z-Vad). P-values were calculated according to the Mann – Whitney U test.

### S9

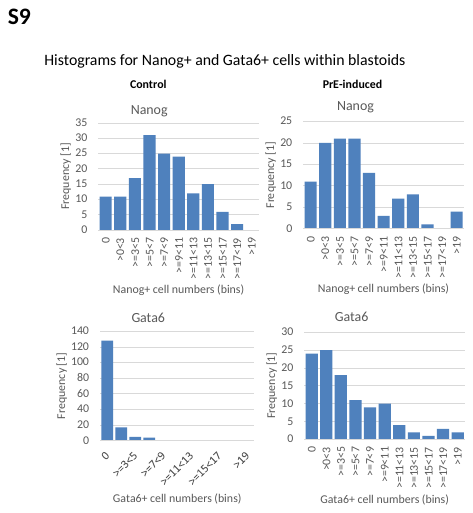

Histograms displaying numbers of either Nanog+ or Gata6+ cells within control and PrE-induced blastoids.
